# diCal-IBD: demography-aware inference of identity-by-descent tracts in unrelated individuals

**DOI:** 10.1101/005082

**Authors:** Paula Tataru, Jasmine A. Nirody, Yun S. Song

## Abstract

**Summary:** We present a tool, diCal-IBD, for detecting identity-by-descent (IBD) tracts between pairs of genomic sequences. Our method builds on a recent demographic inference method based on the coalescent with recombination, and is able to incorporate demographic information as a prior. Simulation study shows that diCal-IBD has significantly higher recall and precision than that of existing IBD detection methods, while retaining reasonable accuracy for IBD tracts as small as 0.1 cM.

**Availability**: http://sourceforge.net/p/dical-ibd

## 1 INTRODUCTION

The notion of identity-by-descent (IBD) between distantly related individuals is playing an increasing role in a variety of genetic analyses, including association mapping (Browning and Thompson, 2012), inferring past demographic history (Palamara *et al*., 2012; Ralph and Coop, 2013), and detecting signals of natural selection (Albrechtsen *et al*., 2010). Currently there exist several useful methods for detecting IBD tracts. These methods are based on characterizing similar haplotypes [e.g., GERMLINE (Gusev *et al*., 2009)] or considering patterns of linkage disequilibrium [e.g., fastIBD and Refined IBD (Browning and Browning, 2011, 2013)], but they do not explicitly model genealogical relationships between genomic sequences. Here, we present a new IBD detection tool, diCal-IBD, which is based on a well-used genealogical process in population genetics, namely the coalescent with recombination. Another feature that distinguishes our method from previous approaches is that we can incorporate demographic information as a prior.

There seems to be no universally accepted definition of IBD. The definition we adopt is the same as that in Palamara *et al*. (2012) and Ralph and Coop (2013). Specifically, an IBD tract is defined as a maximally contiguous genomic region that is wholly descended from a common ancestor without any recombination occurring within the region. In contrast to other methods, we allow IBD tracts to contain point mutations, which are likely to occur in humans due to comparable mutation and recombination rates.

diCal-IBD is able to detect IBD tracts with high accuracy in unrelated individuals, between whom the vast majority of shared tracts are below 1 cM. Existing methods are successful in detecting tracts longer than 2 cM, but have low power for shorter tracts, whereas diCal-IBD maintains reasonable accuracy for tracts as small as 0.1 cM.

## 2 METHOD

diCal-IBD utilizes a recently developed demographic inference method called diCal (Sheehan *et al*., 2013). diCal was originally developed for estimating variable effective population sizes, but it is being extended to handle more complex demographic models, incorporating multiple populations, population splits, migration, and admixture. diCal-IBD will be updated in parallel with diCal and hence will be able to use a complex demographic model as a prior.

diCal is formulated as a hidden Markov model, a decoding of which returns the time to the most recent common ancestor (TMRCA) for each site when analyzing only a pair of sequences. A change in TMRCA requires a recombination event and diCal-IBD uses the posterior decoding of TMRCA to call IBD tracts above a user-specified length, optionally trimming the ends of the tracts that have low posterior probabilities.

diCal requires discretizing time by partitioning it into non-overlapping intervals. The user has the option of specifying any discretization scheme. The default setting implemented in diCal-IBD distributes the pair-wise coalescence probability uniformly over the intervals, similarly as in PSMC (Li and Durbin, 2011), under a constant population size model. An alternative scheme concentrates the intervals in the time period that is most likely to give rise to tracts that are long enough to be detected accurately. This is done by uniformly distributing the coalescence probability conditioned on a random locus being spanned by a tract longer than a user-specified length. Given a variable population size history, we approximate it with a piecewise constant population size.

As an application of IBD prediction, we provide a framework for detecting natural selection. Using the average IBD sharing and posterior probability along the sequence, diCal-IBD identifies regions which exhibit high sharing relative to the background average, indicating possible influence of positive selection.

We refer the reader to the online Supplementary Information for details on data processing, options used in calling diCal, post-processing of posterior decoding, and identification of selection.

## 3 IMPLEMENTATION

diCal-IBD is written in Python 2.7, is platform independent, and has a command line interface which allows the user to completely specify its behavior. The implementation allows for parallel runs of diCal on different sequence pairs. diCal-IBD provides a visualization of the predicted tracts, their posterior probabilities and the corresponding TMRCAs, and sequence-wide average IBD sharing and posterior probability. Accuracy information is also provided if the true IBD tracts are known.

## 4 PERFORMANCE

We carried out a simulation study to compare diCal-IBD with the state-of-the-art IBD detection methods. We used the program ms (Hudson, 2002) to simulate full ancestral recombination graphs (ARGs) for 50 sequences of 10 Mb each. We used a constant recombination rate of 10^-8^, and the African and European demographic histories inferred by Tennessen *et al*. (2012). We simulated sequence data on the ARGs with a constant mutation rate of 1.25 × 10^-8^ per base per generation. From the simulated ARGs, we reconstructed the true pairwise IBD tracts by finding maximally consecutive sites that have the same TMRCA for the pair in question. We only considered the tracts of length *>* 0.1 cM.

To run GERMLINE, fastIBD and Refined IBD, we generated SNP data with approximately 1 marker per 0.2 kb. We assumed perfectly phased data. For further details on running the programs, see Supplementary Information.

Table 1 shows the recall (percentage of true tracts which were correctly recovered), precision (percentage of predicted tracts which were correctly predicted), and F-score (harmonic mean of recall and precision) for each method. See Supplementary Information for other measures of accuracy as a function of the true tract length, as well as the effects of errors in the data, demography, discretization, and trimming based on posterior probabilities. As the table shows, diCal-IBD recalls significantly more tracts with greater precision than other methods, leading to a much higher F-score.

**Table 1.**
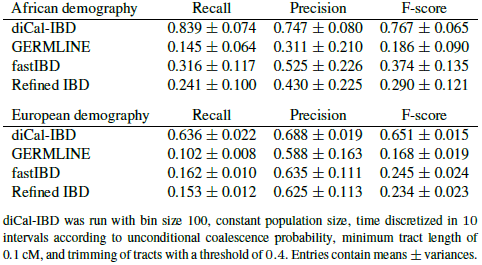
Comparison of IBD detection accuracy.

diCal-IBD was run assuming a constant population size, but its accuracy performance for the examples considered did not seem to be affected much by using this incorrect prior. This suggests that the posterior distribution inferred by diCal is robust to mis-specification of population sizes; whether this trend persists for more complex demographies deserves further investigation.

Figure 1 illustrates the potential of applying diCal-IBD to identify regions under positive selection. We refer the reader to Supplementary Information for further details.

**Fig. 1.**
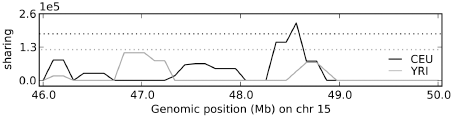
Detection of high sharing using diCal-IBD, on a 4Mb genomic segment (46.0 Mb – 50.0 Mb) on chromosome 15 from Complete Genomics data (Drmanac *et al*., 2010). This region contains a gene thought to be under positive selection in the European population (CEU), located at 48.41 – 48.43 Mb, corresponding with the observed peak in the plot. Dotted lines indicate the thresholds for considering that a region exhibits high sharing.

## ACKNOWLEDGMENTS

We thank Sara Sheehan, Jack Kamm, Matthias Steinrücken, and other members of the Song group for helpful discussions.

## 1 Running diCal

### 1.1 Time discretization

diCal (Sheehan et al., 2013) is formulated as a hidden Markov model based on the coalescent with recombination. For a sample of size 2, hidden states represent the time to the most recent common ancestor (TMRCA) scaled relative to a given haploid reference population size 2*N*_ref_. To apply the standard HMM algorithms, we discretize time into *d* non-overlapping intervals *t*_0_ = 0 < *t*_1_ < ⋯ < *t*_*d*_ = *∞*, with *t*_*i*_ = *g*_*i*_/(2*N*_ref_), where *g*_*i*_ is the number of generations back in time. In diCal-IBD, the user has the option of specifying any discretization scheme. Additionally, we offer two discretization procedures.

Consider the coalescent process in discrete time, with a known population size history, where *N*(*j*) is the size at generation *j*. The probability of two sequences coalescing at generation *g*—that is, the probability that the TMRCA in generations is *g*—is given by 

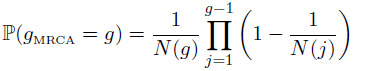

The probability *P*_*i*_ of the TMRCA in generations being placed in a specific interval [*g*_*i*_, *g*_*i*+1_) is 

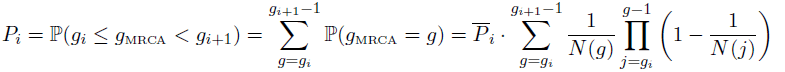

where 

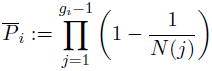

denotes the probability of the TMRCA in generations being older than *g*_*i*_ – 1.

#### Unconditional discretization

This discretization procedure distributes the coalescence probability uniformly over the time intervals by setting 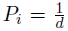 for all *i* = 0*, ⋯, d* – 1. The corresponding values *g*_*i*_ can be determined numerically for increasing values of *i*. When assuming a constant population size, this discretization is similar to the one used in PSMC (Li and Durbin, 2011). The main difference arises from the last time interval. In PSMC, *t*_*d*_ is chosen manually, while here *t*_*d*_ = *∞*.

#### Conditional discretization

The goal of this discretization procedure is to have more time intervals in the period where the TMRCA in generations of tracts longer than *m* cM are expected to be found. Such a discretization scheme should be useful when one is interested in tracts above a certain length while very short tracts (tracts with large TMRCA) are disregarded. To implement this procedure, we need to calculate the coalescence probability at a random locus conditioned on being spanned by a tract longer than *m* cM. Define the following quantities:

- *q*(*g, l*), the joint density that a random locus has TMRCA in generations *g* and is spanned by a tract of length *l*;
- *p*(*g, m*), the joint probability that a random locus has TMRCA in generations *g* and is spanned by a tract of length at least *m*;
- *p*(*m*), the probability of a random locus being spanned by a tract of length at least *m*.

These quantities are related by 

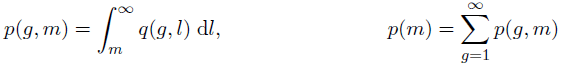

Palamara et al. (2012) showed 

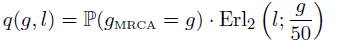

from which we obtain 

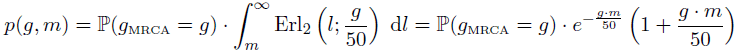

We discretize the time such that 

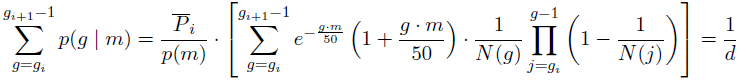

As before, we numerically solve for *g*_*i*_ for increasing values of *i*.

### 1.2 Population size approximation

If the user specifies an arbitrary population size history, diCal-IBD approximates it by a piecewise constant function and uses the approximation in running diCal. The approximation preserves the coalescence probability within each time interval. Let 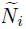 be the approximation of the population size for the time interval [*g*_*i*_, *g*_*i*+1_). Then, the coalescence probability in interval *i* is given by 

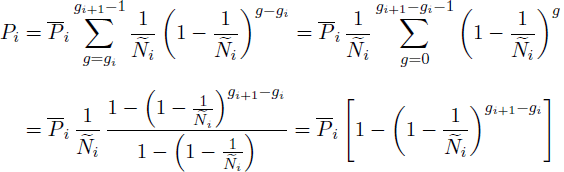

which yields 

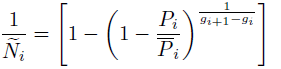

where *P*_*i*_ and 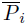 are calculated using the given true population size history.

A constant population size is assumed if no demographic information is provided. Specifically, if the per-generation per-site mutation rate *µ* for a locus is known and the locus consists of *l* sites, diCal-IBD uses *N*(*j*) = *N*_W_ for *j >* 0, where 

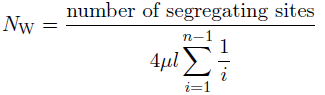

### 1.3 Data binning

As in PSMC (Li and Durbin, 2011), for speed and memory considerations, we group consecutive sites using a user-specified bin size prior to calling diCal. We allow for presence of missing nucleotides (marked as ‘N’) in the data. A bin is marked as ‘missing’ if more than 90% of the nucleotides in the bin are missing, as heterozygous if the bin is not ‘missing’ and at least one base within the bin is heterozygous, and homozygous otherwise. For the analysis described here, we used a bin size of 100.

### 1.4 Other options

When running diCal, we assume that the mutation and recombination rates are given. Before passing on the rates provided by the user, diCal-IBD scales them by multiplying with 4*N*_ref_. We note that due to binning of the input sequences, the mutation and recombination rates are also adjusted by multiplying by the bin size.

Missing heterozygotes (false negatives) can occur due to insufficient read depth. If provided a false negative rate, diCal-IBD lowers the mutation rate to account for missing heterozygotes, as in PSMC (Li and Durbin, 2011).

In addition to mutation and recombination rates, diCal also requires a mutation matrix. Our binning method results in only two types of bins in the data, which we mark as “A” for homozygous bins and “C” for heterozygous bins. Therefore in the mutation matrix, all rates are set to 0, except *A ↔ C*, which are set to 1.

Together with the required input to diCal (the fasta file, using command option -F and the parameters file, using command option -I), we provide the previously described discretization and population size history. As we do not require estimation of the history, we set the number of EM iterations to 0 (using the command option -N 0), the pattern of parameters spanning the time intervals to 1 + 1 + *· · ·* + 1, where the number of 1’s corresponds to a given *d* (using the command option -p), and request the posterior decoding (using the command option -d 5).

For the presented analysis we ran diCal v1.2.

## 2 Processing diCal output

### 2.1 Identification of tracts

We call IBD tracts using the posterior decoding from diCal, as maximally consecutive sites that have the same TMRCA for the pair in question. We disregard the tracts for which the TMRCA is placed in the last time interval, as this could be an artifact resulting from the lack of finer intervals. Provided that there are enough time intervals, this removal should not change the performance of diCal-IBD drastically, as tracts with old TMRCA are expected to be very short.

As we run diCal on binned data, we recalculate the position of the detected tracts to correspond to the original sequence length.

For reporting and accuracy calculation purposes, we only consider tracts above a user-defined length. For the presented analysis, we used a minimum length of 0.1 cM.

### 2.2 Trimming of tracts

In addition to the inferred TMRCA for each predicted tract, we also obtain the corresponding posterior probabilities, which can be used as a measure of confidence for our prediction along the tract. We optionally use this extra information by discarding bins and tracts whose posterior probabilities are too low. To trim a tract, we discount bins at both ends if the corresponding posterior probability is below a given threshold. After trimming, we keep the tract only if the average posterior probability of the remaining bins is above the threshold. Reporting of tracts is contingent only on the original length; that is, we still report trimmed tracts even if they are shorter than the specified minimum length. However, for our accuracy calculations we do not count the contribution of tracts which do not meet the minimum length requirement after trimming.

## 3 Simulations

### 3.1 Simulation of ARGs

We used the program ms (Hudson, 2002) to simulate full ancestral recombination graphs (ARGs) with a per-base recombination rate of 10^-8^ for 50 sequences of 10 Mb each. We used two different population histories, corresponding to the African (AF) and European (EA) demographic histories inferred by Tennessen et al. (2012). As shown in Figure 1a, both histories contain an ancient bottleneck and periods of rapid population expansion in the recent past. The European population contains the out-of Africa bottleneck and a more recent bottleneck and, as well as two different epochs of rapid expansion.

**Figure 1:**
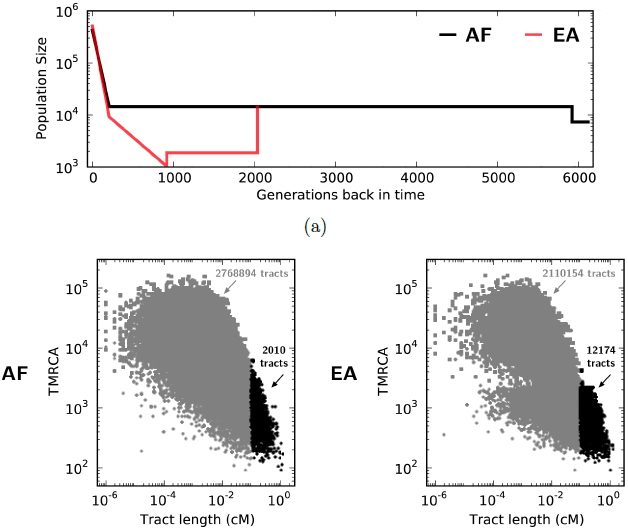
Population histories used for simulating data: African (AF) and European (EA) as inferred by Tennessen et al. (2012). (a) Population size histories used. AF and EA share a common history until almost 2000 generations ago. (b) Simulated pairwise tracts as a function of their length and TMRCA in generations. The black dots correspond to the tracts longer than 0.1 cM, which are used in the presented analysis, out of which only 4 (AF) and 5 (EA) are above 1 cM.

To calculate the scaled recombination rate and the corresponding population sizes, we used an *N*_ref_ of 1861. The ms commands used for simulating the ARGs for the AF and EA demographies are

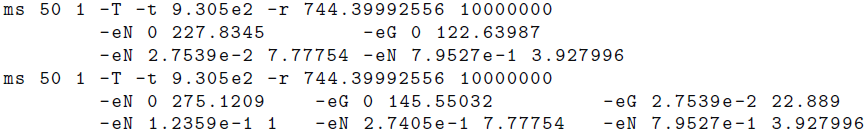

We note that the -t option is only used for later recovery of the mutation rate.

Figure 1b shows scatter plots of the simulated tracts. Due to the low number of simulated sequences, very few tracts are above 1 cM. The higher number of tracts above 0.1 cM in the EA dataset is a result of the reduced population size after the out-of-Africa bottleneck. This event left a strong signal in the data, creating a clear separation between the tracts that are older and the ones that are younger than the bottleneck.

### 3.2 Simulation of sequences

We generated sequence data on the ARGs using a per-base mutation rate of 1.25 *×* 10^-8^ (Kong et al., 2010, 2012). In real data, errors in base calling can lead to non-existent variants being called (false positives) or to true variants being overlooked (false negatives). To account for this, we superimposed sequencing errors, adding variants with rate 6 *×* 10^-6^ and removing variants with rate 0.02, in accordance with reported false positive and false negative rates (Illumina, 2011) and previous studies (Browning and Browning, 2013). To remove variants, we paired the sequences (to mimic diploids) and, for each heterozygous position, we changed one of the two alleles so that the site became homozygous. Due to our pairing of the sequence, we added errors with probability twice the false negative rate. To add false positives, we chose the number of false SNPs from a Poisson distribution with mean equal to the false positive rate *×* sequence length *×* number of sequences. These additional SNPs were then distributed uniformly over the sequences.

To run other programs, we generated SNP data from the simulated sequence according to the SNP density (15 million SNPs genome wide) reported by the 1000 Genomes Project Consortium (2010). After inclusion of all segregating sites, we randomly selected additional sites such that the resulting marker density is approximately 1 marker per 0.2kb.

Table 1 shows the summary of the simulated sequence data. The removed variants were not necessarily from different SNPs, so the total number of SNPs affected by the false negative errors is lower than the number of removed variants. Some of the SNPs could be present in very low frequency, so a removed variant could lead to the complete loss of the SNP. The EA dataset had an overall lower number of SNPs, most likely a consequence of the out-of-Africa bottleneck.

**Table 1:** Summary of the simulated sequences under the two population histories.

|  | Simulated SNPs | Removed variants | Affected SNPs | Removed SNPs | Added SNPs |
| --- | --- | --- | --- | --- | --- |
| AF | 24370 | 4368 | 3809 | 164 | 3059 |
| EA | 12678 | 3088 | 2624 | 80 | 3077 |

## 4 Accuracy measures

From the simulated ARGs, we reconstructed the true pairwise IBD tracts by finding maximally consecutive sites that have the same TMRCA for the pair in question. We note that this excludes recombinations which do not affect the coalescence time, an event which occurs only rarely. We only considered the tracts of length *>* 0.1 cM.

To calculate the accuracy, we use the percentage overlap between true and predicted tracts, rather than their lengths. Doing so, all tracts, regardless of their length, contribute equally to the accuracy measures. We require a minimum percentage overlap *a* (here set to 0.5) for a prediction to be considered as a true positive. Let true_*i*_ be a true tract, pred_*i*\*_ be its best prediction (the predicted tract with highest overlap) and pred_*j*_ be a predicted tract. The accuracy measures that we calculate are

*•* true positive: overlap between true_*i*_ and pred_*i*\*_ if the percent overlap (with respect to the length of both the true and the predicted tract) is greater than *a*;
*•* false negative: any part of true_*i*_ that does not overlap with any predicted tract;
*•* false positive: any part of pred_*j*_ that does not overlap with any true tract;
*•* power, all overlaps between true and predicted tracts;
*•* under-prediction: the part of true_*i*_ that does not overlap with pred_*i*\*_;
*•* over-prediction: the part of pred_*i*\*_ that does not overlap with true_*i*_.

All measures listed above are calculated as a function of the length of the true tract, except the false positive, which is a function of the length of the predicted tract. In addition to the above measures, we report recall (the percentage of true tracts which were correctly recovered), precision (the percentage of predicted tracts which were correctly predicted), and F-score (the harmonic mean of recall and precision). Together, these provide an overall measure of accuracy.

Let 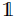 be the indicator function. Then we can define mathematically the accuracy measures as follows. We note that true*i* represents both the tract and its length.

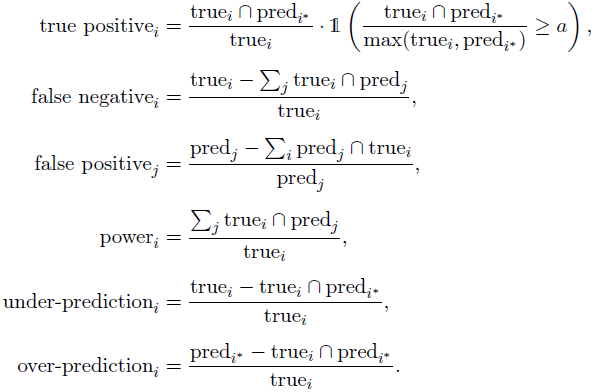

Figure 2 gives an overview of the different types of overlaps. Both true and predicted tracts are numbered, while for the predicted tract, the corresponding *i*^*^ index is given after the equal sign.

**Figure 2:**
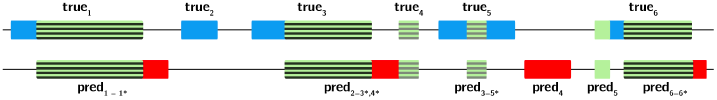
Types of overlaps. Regions in both true and predicted tracts are coloured such that: green indicates overlap between true and predicted tracts and blue (red) indicate regions in true (predicted) tracts that to not overlap with other predicted (true) tracts. Horizontal lines (both gray and black) mark, for each true tract, the best overlap with a predicted tract. Black horizontal lines indicate that the overlap in percentage is above the required threshold *a*. For details, see main text.

A predicted tract can be the best prediction for several true tracts. The overall average accuracy for the example in the figure is then given by 

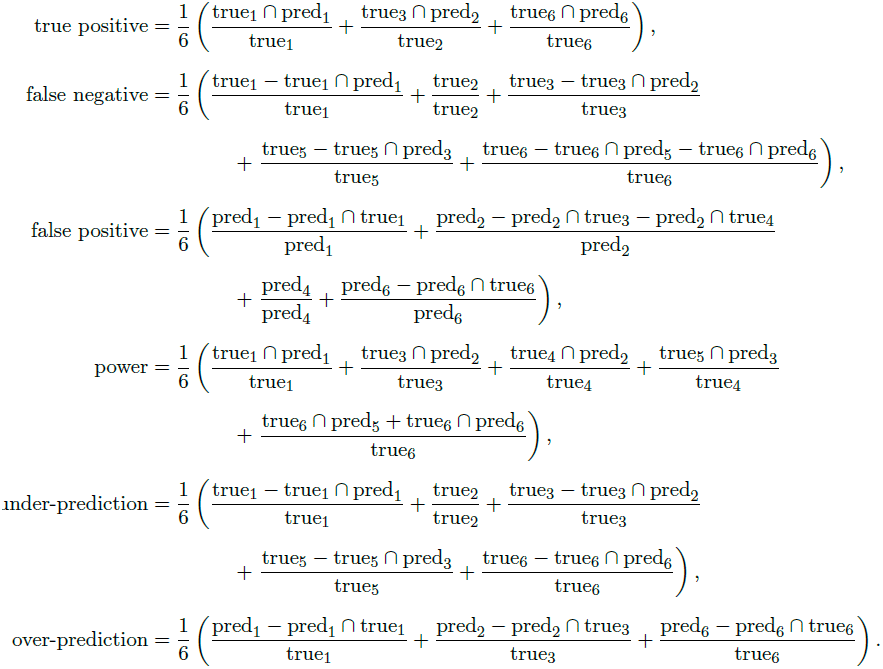

## 5 Detecting selection

Positive selection has been shown to result in an increase in IBD sharing (Albrechtsen et al., 2010; Han and Abney, 2013). We provide a framework within diCal-IBD to identify genomic regions with high sharing relative to the background average. We calculate the sharing for non-overlapping windows as the total length of the tracts spanning each window, divided by the number of pairwise comparisons. We consider a window to have increased sharing if it is greater than three standard deviations from the average sharing across all sites. Similarly, we calculate the average posterior probability by considering the total average posterior of the tracts spanning the windows, divided by the number of tracts per window. In the presented analysis we used a window size of 0.1 cM.

## 6 diCal-IBD implementation

diCal-IBD is implemented in Python 2.7, using Python packages by Hunter (2007); Jones et al. (2001); Mailund (2006). diCal-IBD is freely available at http://sourceforge.net/projects/ dical-ibd and is accompanied by an instruction manual.

## 7 Running other programs

We compare the performance of diCal-IBD on the simulated data with three IBD detection softwares

- GERMLINE version 1.5.1 (Gusev et al., 2009);
- fastIBD, packaged with BEAGLE version 3.3.1 (Browning and Browning, 2011);
- Refined IBD, packaged with BEAGLE version 4 (Browning and Browning, 2013).

In the following we describe the options we used while running these programs. We note that this improved the performance compared to the default settings. Only GERMLINE allows for correcting errors in the data. To create diploid input for these programs, we replicated and paired together each sequence to generate “diploid” data which was homozygous at all sites.

### 7.1 GERMLINE

As we only considered IBD segments larger than 0.1 cM, we set **min_m** = 0.1. We also made use of GERMLINE’s **h_extend** option, which improves performance when data is well-phased. The **bits** parameter determines the number of markers to be considered in an exact matching seed. Choosing a value which is too large may result in missing shorter IBD segments, while one that is too small affects computation time. We set this parameter so that a seed length corresponded to 0.02 cM, which was significantly shorter than our threshold for candidate IBD segments (0.1 cM). For example, for a density of 1 marker per 0.2kb, 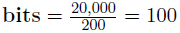. Note that the recombination rate we used in the simulated data, *r* = 10^-8^, implies that 0.02 cM corresponds to 20kb. When running on perfect data, we set the allowed number of homozygous mismatches per seed **err_hom** to 0. Otherwise, we calculate the number of expected errors in a bin (here, 20kb). Using this method, we set **err_hom** = 1. Because our data consisted of haploid sequences, we set **err_het** = 0.

### 7.2 fastIBD

We ran fastIBD with the default setting **ibdscale** = 2. Because of no phase uncertainty in our data, we ran only one iteration (**ninterations=1**) and allowed for a very high score threshold of **fastibdthreshold**= 10^-2^.

### 7.3 Refined IBD

Refined IBD uses the GERMLINE algorithm to find candidate IBD segments, and the **ibd-window** parameter is equivalent to the GERMLINE **bits** parameter, and was set to the same value (**ibd-window** = 100). Likewise, we set **ibdcm** = 0.1, as this parameter is equivalent to GERMLINE’s **min_m**. Due to no phase uncertainty in our data, we set the threshold for the LOD score (the base 10 log of the likelihood ratio) to **ibdlod** = 1.0.

## 8 Results and discussion

Figures 3 to 5 show the performance of GERMLINE, fastIBD, Refined IBD and diCal-IBD for the two simulated datasets. From the figures, it is clear that diCal-IBD has an overall better performance, with an increased recall, while the precision is at least as high as for the other programs. Overall, all programs have better accuracy for the AF dataset. We believe this is mainly due to the higher number of SNPs present in the AF dataset (Table 1), which increases the amount of available information.

**Figure 3:**
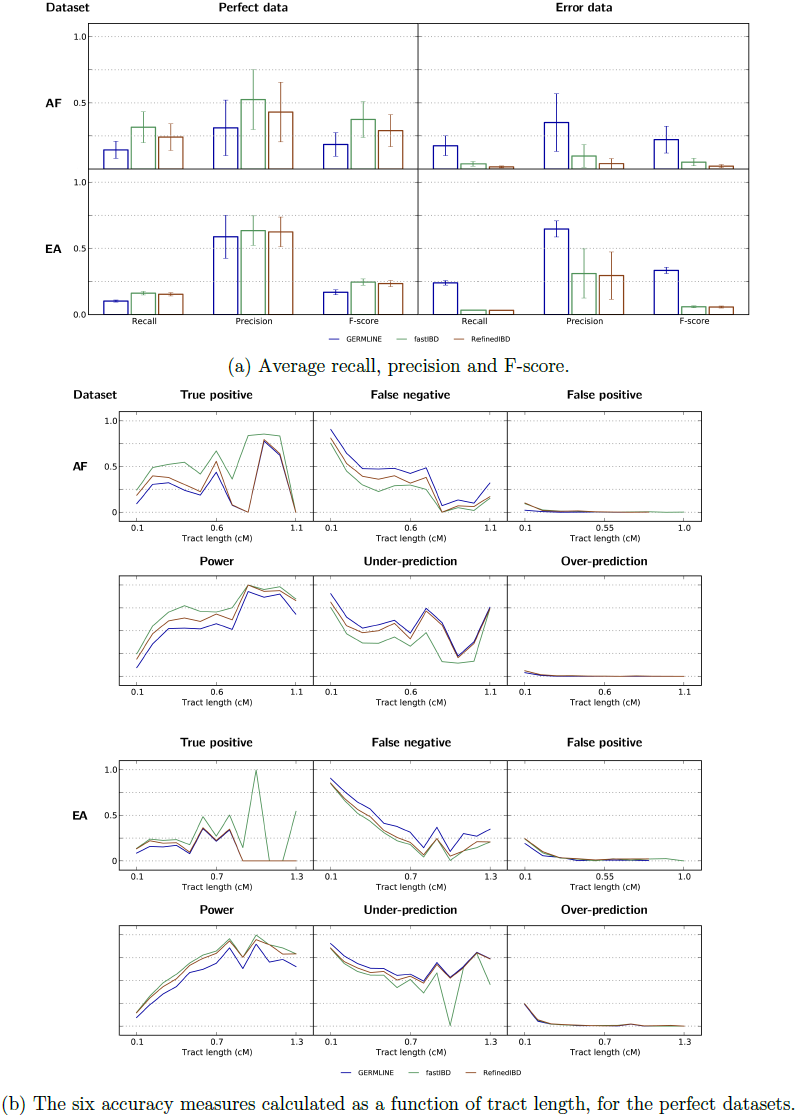
Accuracy results for GERMLINE, fastIBD and Refined IBD for both AF and EA datasets.

**Figure 4:**
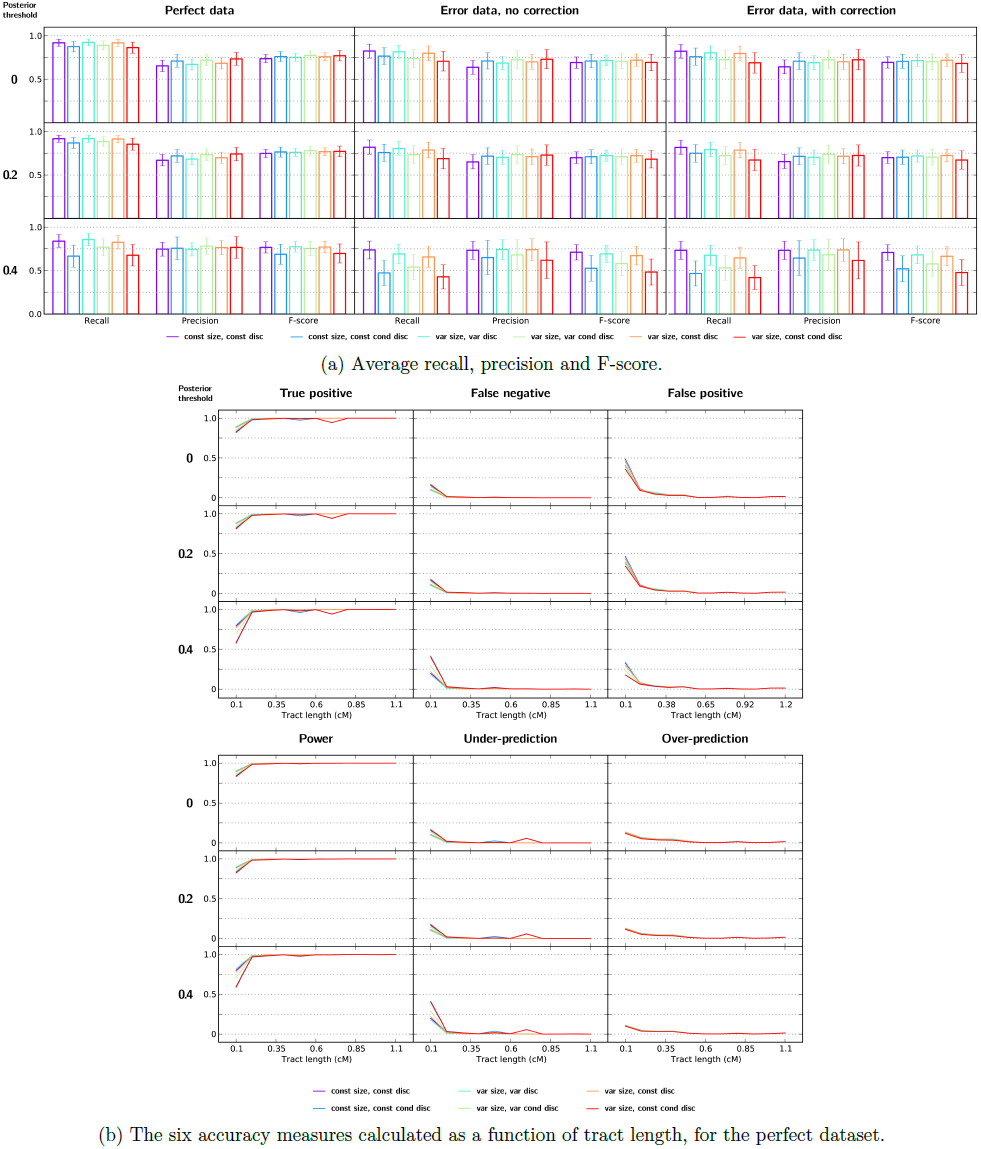
Accuracy results for diCal-IBD, using different population sizes, discretizations and posterior probability thresholds for trimming of tracts, for the AF dataset.

**Figure 5:**
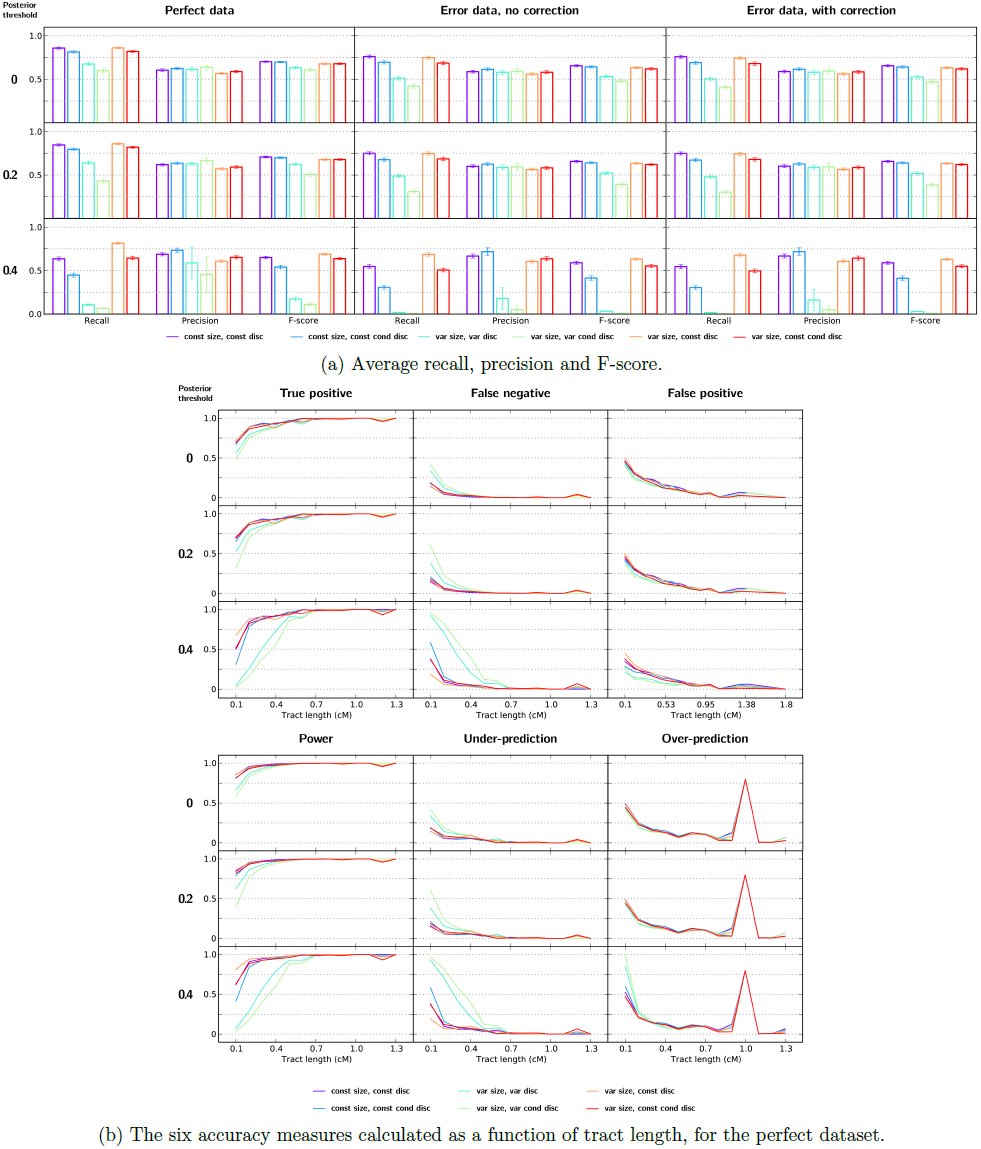
Accuracy results for diCal-IBD, using different population sizes, discretizations and posterior probability thresholds for trimming of tracts, for the EA dataset.

In Figures 3b to 5b, we plotted the accuracy as a function of tract length using windows of cM (i.e., the first window contains tracts between 0.1 cM and 0.2 cM in length, the next 0.2 cM – 0.3 cM, and so on). We note that the sudden jumps in the figures for the longer tracts are due to the low number of tracts in those ranges (Figure 1b). diCal-IBD has very high true positive and power for tracts as low as 0.1 cM, while the other programs recover only a modest portion of the tracts. The small difference between the true positive and power for diCal-IBD indicates that most true tracts overlap at least 50% with a predicted tract. This, together with the similarity between false negative and under-prediction, suggests that most often there is a unique correspondence between true and predicted tracts, in that the situations where one true tract overlaps with several predicted tracts, or vice versa, are rare.

The main fault in diCal-IBD is its relatively high false positive rate. For the AF dataset, the discrepancy between the false positive and over-prediction rates implies that a significant portion of the false positives correspond to predicted tracts that do not overlap with any true tract. These tracts may correspond to true tracts too short to pass the length filter; this is supported by the fact that the false positive rate is considerably higher in the first window considered. In contrast, for the EA dataset, the false positive and over-prediction rates are more comparable. This indicates that false positives in this prediction are the result of extension at the boundaries of true tracts.

### Effect of errors in the data

Sequencing errors in the data reduce diCal-IBD’s accuracy slightly, with a more pronounced effect for the EA dataset. Correcting for the false negative errors doesn’t seem to affect the results. In contrast, the other programs seem to be more drastically affected by errors. GERMLINE shows a better performance for the datasets with errors. We believe this is a consequence of GERMLINE’s option to accommodate mismatches in tracts (the **err_hom** parameter, which was set to 1), which effectively allows for presence of point mutations inside tracts. fastIBD and Refined IBD do not allow for mismatches. While this option in GERMLINE could be used to account for point mutations even in perfect datasets, the number of such changes is highly dependent on the population history.

### Bin size

In the presented analysis we binned the sequences using a bin size of 100. To investigate whether this influences the performance, we reran diCal using a bin size of 10. The performance showed no difference between the two runs (results not shown).

### Population size and discretization

We ran diCal-IBD assuming a constant population size (‘const size’) or approximating the true population size (‘var size’). For the former, we used the two discretizations described previously (‘const disc’ and ‘const cond disc’, respectively), considering tracts longer than *m* = 0.01 cM for the conditional discretization. For the latter, we recalculated the two presented discretizations under the variable population size (‘var disc’ and ‘var cond disc’), but also used the two discretizations under a constant size. This enables us to investigate the effect of using different population sizes when the discretization is fixed.

diCal-IBD generally shows similar F-scores under the different assumed population sizes and discretizations, indicating the robustness of the posterior distribution. The discretization seems to have a higher impact on the performance, as the recall and precision vary with the discretization used, but not with the population size assumed.

#### Trimming of tracts

We investigated the effect of trimming of tracts based on the posterior probability, using thresholds 0.2 and 0.4. It is clear that in most cases the precision increases, while the false positive and over-prediction are reduced. Trimming can, however, have the opposite desired effect, as it potentially removes ends of the predicted tracts that overlap correctly with a true tract. Most of the effects of trimming are rather minor, indicating that tracts have overall posterior probabilities above 0.4. In some cases, trimming using 0.4 as threshold decreases the recall more drastically, without a large enough increase in the precision, to balance the final F-score.

#### Density of time intervals

From our experiments, we observed that using time intervals that are too dense decreases the performance of diCal-IBD. This is somewhat counter-intuitive, as one would expect that using more time intervals would allow diCal to identify the TMRCA of each tract more accurately. However, we believe the decline in performance to be the result of the large variation in TMRCA for tracts of the same length (Figure 1b). On the other hand, discretization schemes that are too sparse can blur the boundaries between adjacent IBD tracts. An increased density of time intervals can arise from using a conditional discretization with a high tract length threshold tracts (for example, 0.1 cM), or from the discretizations based on the variable population size. Such an example is the discretization for the EA dataset based on the variable population size, where the resulting dense intervals are probably due to the out-of-Africa bottleneck. The low confidence in the TMRCA is also reflected in the effect of trimming of tracts, as the performance is drastically affected when using a threshold of 0.4.

### 8.1 Detecting selection

For illustrating the visualization method for identifying regions under positive selection, we used a 4 Mb segment of chromosome 15 (46 Mb – 50 Mb) from the 69 Genomes public dataset by Complete Genomics (Drmanac et al., 2010). In this region, the SLC24A5 gene is located. This gene is responsible for light skin pigment and lack of dependence on sunlight for vitamin D production, and has been found to be under positive selection in northern Europeans (Wilde et al., 2014).

We ran diCal-IBD on samples from both European (CEU) and African (YRI) populations (Table 2). We assumed the mutation and recombination rates to be 1.25 *×* 10^-8^ and 10^-8^, respectively. We used constant population size, the unconditional discretization and untrimmed tracts that were at least 0.1 cM long. Figure 6 shows the resulting average sharing and posterior probability. The CEU data contains one peak with high sharing corresponding to the region (48.41 – 48.43 Mb) which has been found under positive selection. As expected, the YRI data does not exhibit high sharing.

**Table 2:** CEU and YRI samples from Complete Genomics data.

| Coriell ID | Population | Coriell ID | Population |
| --- | --- | --- | --- |
| NA06985 | CEU | NA18501 | YRI |
| NA06994 | CEU | NA18502 | YRI |
| NA07357 | CEU | NA18504 | YRI |
| NA10851 | CEU | NA18505 | YRI |
| NA12004 | CEU | NA18508 | YRI |

**Figure 6:**
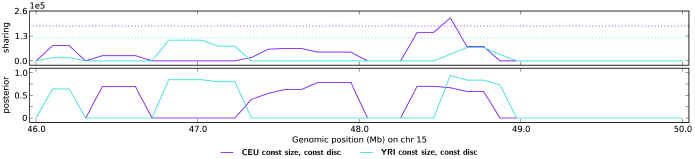
Detection of high sharing using diCal-IBD. Dotted lines indicate the thresholds for considering that a region exhibits high sharing.

